# Comparative pangenomics of *Trypanosoma brucei* reveal host specialization is driven by targeted or segmental genome streamlining in human and equine-restricted lineages

**DOI:** 10.64898/2026.06.15.732286

**Authors:** Muyiwa S. Adegbaju, Olanrewaju B. Morenikeji, Prakash K Singh, Xiomara Lane, Olusola Ojurongbe, Bolaji N. Thomas

**Author notes:** Corresponding Authors; Bolaji N. Thomas, Muyiwa S. Adegbaju.

## Abstract

How the eukaryotic parasite *Trypanosoma brucei* remodels its genome during host specialization remains a fundamental question in evolutionary biology. In contrast to the uniform, systemic gene loss seen in resident bacterial genomes, the structural variations driving niche restriction in extracellular pathogens are poorly defined. Here, we establish a high-resolution comparative structural pangenomic map of the *T. brucei* complex. By analyzing the human-restricted specialist DAL972 and the equine-restricted specialist *Trypanosoma equiperdum* IVM-t1 against the conserved TREU927 backbone, we determine how host-vector constraints dictate eukaryotic genome architecture. Our analyses reveal that host specialization is governed by a coordinated spatial blueprint that segregates structural variation into two distinct evolutionary pathways: targeted perimeter streamlining and systemic core erosion. The human specialist DAL972 preserves its core metabolic toolkit under intense purifying selection, restricting genomic variation to the compartmentalized trimming of its sub-telomeric variant surface glycoprotein (*VSG*) archives. Conversely, the equine specialist IVM-t1, which has completely abandoned cyclical tsetse fly transmission, displays multi-focal core erosion across internal domains. We demonstrate that vector abandonment forces a shift to strict asexual clonal population dynamics. This transition eases purifying selection, allowing obsolete, insect-stage flagellar and cytoskeletal structures to decay through neutral genetic drift. At the chromosomal perimeters, this structural allocation unmasks an unannotated sub-telomeric reservoir of 86 profile-validated retrotransposon hot spot (RHS) and catalytic DDE transposase domains organized into 23 discrete loci. Operating under adaptive positive selection in the human specialist, these silent, repetitive elements function as physical recombination anchors that stabilize strand exchange. This mechanism drives rapid antigenic diversification while insulating the core genome from dangerous macro-rearrangements. Importantly, we bridge this macro-scale chromosomal architecture to micro-scale proteomic execution through tertiary modeling of these unannotated specialized loci. This structural translation unmasks a biophysical mechanism of structural sequestration, where lineage-specific mutations and specificity-determining positions are deeply buried within hydrophobic protein cores to shield functional divergence from host immune surveillance. By pairing these rigid, hidden core packing networks with hyper-variable surface tiles and dynamic boundary hinges, the parasite deploys tiered molecular mimicry to selectively manipulate host cellular networks. Together, these findings redefine our understanding of structural and non-coding architecture as mechanical drivers of parasitic adaptability, uncovering a direct pipeline from pangenomic spatial variations to stable, structure-based allosteric drug targets.

## Introduction

The African trypanosome, *Trypanosoma brucei*, serves as a premier model for evolutionary plasticity, existing as a complex of lineages that range from broad-spectrum generalists to highly host-restricted specialists [1,2]. These protozoan parasites are the causative agents of African trypanosomiasis a suite of devastating diseases traditionally restricted to 36 countries within the sub-Saharan tsetse belt [3]. Within this endemic region, the disease manifests in two distinct forms. Human African Trypanosomiasis (sleeping sickness) is caused by *T. b. gambiense* and *T. b. rhodesiense*, which utilize localized molecular factors such as the *TgsGP* and *SRA* genes to neutralize host innate immune barriers [4–7]. In contrast, Animal African Trypanosomiasis (nagana) is driven by tsetse-dependent subspecies like *T. b. brucei*, severely decimating sub-Saharan livestock populations and causing annual economic losses exceeding $4.5 billion [3,8]. Natively, the pathobiology of the *T. brucei* complex is defined by cyclical transmission via the tsetse fly vector (*Glossina* spp.) and a remarkable system of antigenic variation [9]. Mammalian bloodstream forms evade host adaptive immunity by periodically switching a dense surface coat of variant surface glycoproteins (VSGs) drawn from a massive genomic archive [10,11]. Despite over a century of research, controlling these endemic lineages remains a formidable challenge due to late-stage drug toxicity, emerging resistance, and the frustration of vaccine development by this perpetual antigen-switching capacity [8].

Crucially, the biological boundaries of the *T. brucei* complex extend far beyond these traditionally defined, tsetse-bound subspecies. Vector-independent derivatives of this complex have successfully breached historical geographic and insect-vector constraints [2]. By adapting to mechanical transmission via biting flies or venereal transfer, non-tsetse lineages such as *Trypanosoma evansi* and *Trypanosoma equiperdum*—an equine specialist—have extended the socioeconomic and agricultural burden of this parasite complex globally into expansive territories across Asia and South America. While historically considered distinct species based on clinical presentation and the loss of kinetoplast DNA (dyskinetoplasty), recent genetic analyses confirm that *T. equiperdum* and *T. evansi* represent monophyletic clusters that emerged independently from *T. brucei* ancestors. However, characterizing the extensive genomic remodeling that underwrites this transition to host-restriction has been severely constrained by a historical reliance on a single linear reference genome (TREU927). This inherent reference bias systematically obscures non-reference structural variants including large-scale inversions, segmental duplications, multi-focal core deletions, and hyper-variable sub-telomeric locus dynamics, that drive host adaptation [1,10,12,13]. As genome biology shifts toward a pangenomic paradigm, capturing this full spectrum of structural variation is essential for deciphering how these specialized lineages evolve [13].

The necessity of this study is underscored by a fundamental gap in our understanding of how large-scale genomic architecture influences host restriction. Recent *de novo* assemblies of related trypanosomatids, such as *Trypanosoma congolense*, have demonstrated that while core chromosomal regions remain syntenically stable, sub-telomeric zones serve as dynamic laboratories for antigenic innovation and structural reshaping [14]. Crucially, *T. congolense* lacks the canonical variant surface glycoprotein (VSG) expression sites found in *Trypanosoma brucei*, confirming that different parasitic species deploy highly divergent strategies for immune evasion and genome organization [14]. While it is recognized that host-specialized lineages within the *T. brucei* complex itself exhibit distinct phenotypic traits and genetic losses, the underlying structural rules governing how these independent genomes remodel their functional loci remain poorly defined. Specifically, it remains unclear whether niche restriction across different host-vector interfaces forces these intra-species lineages to follow a uniform trajectory of genome reduction, or if they utilize divergent, spatially organized strategies of structural modification to reshape their functional architecture. Resolving this distinction is critical to determining whether mechanical constraints on chromosome architecture dictate the evolutionary boundaries of parasite adaptability.

This structural reshaping occurs within an evolutionary lineage that is profoundly distant from canonical model organisms. *T. brucei* belongs to the Discoba clade, which shared its last common ancestor with animals, plants, and fungi approximately 1.5 billion years ago. This vast evolutionary divergence presents a significant barrier to traditional functional genomics, as approximately 50% of the *T. brucei* proteome (∼4,900 genes) consists of hypothetical proteins lacking identifiable orthologs in traditional model eukaryotes [15,16]. While high-throughput forward genetic screens have systematically mapped essential cell-cycle phenotypes across these unannotated loci [17], structural characterization was historically limited. Because deep-learning structural prediction frameworks rely on co-evolutionary signals extracted from multiple sequence alignments (MSAs), the lack of heavily sampled sequenced sister taxa resulted in shallow alignment profiles and low-confidence 3D models. However, recent expansions in global eukaryotic and environmental sequence repositories have fundamentally resolved this limitation. By leveraging deeply sampled kinetoplastid and protistan genomic data through optimized alignment search architectures, recent workflows have successfully rescued previously unmodelable profiles [18]. Integrating these expanded evolutionary profiles into high-throughput structural prediction networks such as AlphaFold2 paired with custom and updated MSA generation pipelines now permits high-confidence, routine 3D modeling of complex, trypanosomatid-specific domains.

Our study addresses this challenge by establishing a high-resolution structural pangenomic framework for the *T. brucei* complex. Rather than relying on fragmented, population-level short-read datasets that mask large-scale chromosomal rearrangements, we leverage end-to-end, chromosome-level *de novo* assemblies of the human specialist (DAL972) and the equine specialist (*T. equiperdum* IVM-t1) mapped directly against the conserved TREU927 reference backbone. This architectural approach allows us to track continuous synteny and spatial variations across entire chromosomes, resolving this ambiguity by demonstrating that host specialization is governed by a highly segregated spatial blueprint. By integrating macro-scale chromosomal topography with micro-scale tertiary structural modeling, this study charts the specific genetic boundaries and directional selective pressures that govern host-restricted niche isolation.

Crucially, by resolving the 3D atomic architecture of previously unannotated, lineage-specific proteins within highly reorganized sub-telomeric regions, this work unmasks how structural sequestration hides functional divergence from host immunity. Ultimately, this multi-scale structural blueprint shifts the paradigm of parasite evolutionary genomics from descriptive gene-counting to a mechanism-driven pipeline for structure-based, allosteric drug discovery.

## Materials and Methods

### 3.1 Data acquisition and assembly quality control

High-fidelity, chromosome-level genome assemblies for four distinct *Trypanosoma brucei* complex (subgenus *Trypanozoon*) strains were retrieved from the NCBI BioProject repository under the following accession numbers: PRJNA11756 (*T. b. brucei* TREU927 genomic reference anchor), PRJEA40697 (*T. b. gambiense* DAL972 human-restricted specialist), PRJNA477427 (*T. equiperdum* IVM-t1 equine-restricted specialist), and PRJNA723622 (*T. b. brucei* EATRO1125 livestock-derived generalist anchor). Selection was strictly limited to assemblies within the *Trypanozoon* subgenus possessing fully resolved, telomere-to-telomere megabase chromosomal scaffolds to eliminate technical assembly gaps as confounding variables in downstream pangenomic comparisons. Assembly integrity and gene-set completeness were systematically verified using BUSCO (v5.8.3) executed against the euglenozoa_odb10 baseline database [20]. Macro-scale topology, N50 metrics, and contiguity profiles were audited via QUAST (v5.3.0), focusing exclusively on the eleven megabase-sized core chromosomes [21]. To safeguard against coding-density anomalies introduced by unplaced short scaffolds, a normalized assembly index was calculated for each strain, defined as the ratio of total core megabase chromosome length (Mb) to total predicted coding sequences (CDS) extracted from corresponding NCBI structural annotations [22–24].

### 3.2 Pangenomic intersection modeling and quality filtering

Orthologous gene groups (orthogroups) and unassigned singletons were delineated across the retrieved genomes using OrthoFinder (v2.5.4) [22,23]. To prevent sequence fragmentation from corrupting downstream sliding-window density calculations and evolutionary selection () models, the highly fragmented *T. brucei* EATRO1125 assembly (>700 contigs) was excluded from primary macro-structural alignment matrices. Because unplaced contig boundaries introduce artificial sequence gaps that inflate non-synonymous substitution rates (*d_N_*). However, EATRO1125 was utilized exclusively as an independent, un-scaffolded validation database to verify core metabolic deletions and sub-telomeric loci via targeted ortholog reconciliation. The remaining high-contiguity, chromosome-level assemblies i.e the TREU927 reference anchor (7,777 loci), the DAL972 human specialist (7,614 loci), and the IVM-t1 equine specialist (6,460 loci), were partitioned into a tripartite structural pangenome matrix within the R statistical environment. Intersection vectors programmatically classified orthogroups into three mutually exclusive categories: core compartments symmetrically conserved across all three assemblies, accessory compartments shared by exactly two strains, and unique compartments restricted to a single specialized lineage. Geometric rendering of these pangenomic intersections was executed using the VennDiagram package (v1.7.3) under unscaled parameters to prioritize the clear visualization of shared and unique sectors [25].

### 3.3 Spatial matrix generation and chromosomal segmentation

To establish a spatially continuous evolutionary map, the compiled pangenomic classifications (7,966 total orthogroups) were integrated into a unified spatial matrix mapped sequentially along the primary axes of the TREU927 reference genome coordinate anchor. Absolute numeric indices were assigned to each orthogroup based on sequential reference locus tag suffixes. Loci were partitioned into three continuous architectural tracks using a tripartite Boolean filter: the stable core framework, segmental erosion footprints (loci absent in IVM-t1), and variable profiles (lineage-isolated sequences unique to DAL972). To resolve this master spatial matrix into discrete chromosomal data frames, an automated segmentation script was applied to isolate the eleven canonical megabase chromosomes based on NCBI chromosome accession boundaries. For each segmented chromosome, total reference gene capacities and regional track occupancies were quantified. Localized track counts were normalized relative to total chromosomal gene volumes to generate a standardized genomic profile matrix for cross-chromosomal comparison.

### 3.4 Sliding-window density analysis and boundary delineation

Architectural transition zones between stable chromosomal cores and variable sub-telomeric provinces were delineated using a sliding-window algorithm deployed across the sorted reference gene index axis of the master spatial matrix. The sliding window utilized a fixed aperture of 50 consecutive gene loci with a step-size of 10 loci. Within each window, local densities were quantified for the three pangenomic tracks defined in Section 3.3. Physical coordinates for each window were assigned based on the median gene index of the 50-loci block to ensure geometric centering. To pinpoint the precise structural boundaries where the genome stabilizes into the core assembly, an empirical filter was applied to the density tracks: the outer boundary of the sub-telomeric zone was mathematically defined as the final coordinate hotspot where regional segmental erosion footprints (loci absent in IVM-t1) dropped and remained consistently below a 2% occupancy baseline. Features residing distal to these transition zones within the defined sub-telomeric compartments were isolated as a raw pool of 2,207 unannotated loci, which were subsequently subjected to selective pressure filtering and tertiary structural modeling.

### 3.5 Statistical validation and radial circlize mapping

To determine whether genomic remodeling across specialized lineages followed a uniform pattern of stochastic decay or was driving regional, non-uniform structural trends, a chromosome-wide statistical census was executed. Observed structural variants (SVs) comprising segmental erosion in IVM-t1 and gene loss in DAL972 were aggregated across the eleven canonical chromosomes. An expected null-hypothesis distribution matrix was generated where the expected rate of sequence alteration for each chromosome was calculated as a strict proportion of its gene capacity relative to the global genome backbone. Departures from size-proportional expectations were evaluated using a Pearson’s Chi-Square goodness-of-fit test implemented via the scipy.stats Python package [26]. This macro-scale validation established a statistically significant non-uniform distribution of structural modifications, justifying localized downstream adaptive selection profiling. The global distribution and regional density of these lineage-specific modifications were mapped using a radial visualization framework via the circlize package within the R statistical environment [27]. Structural variant intervals were imported in standardized BED format and mapped to Chromosomes 1–11, filtering out unplaced scaffolds. Spatial divergence scattering was assessed through a dual-track analytical layer. The primary track comprised an inter-variant distance rainfall plot using log10-transformed physical distances between consecutive deletion breakpoints to resolve regional clustering. Concurrently, regional variant density was quantified on a secondary track using a rolling heatmap calculated via a sliding-window function with a fixed window of 50 kb. Radial sectors were initialized with explicit spacing at chromosomal boundaries, and coordinate scales were standardized across all tracks relative to the TREU927 reference backbone.

### 3.6 Locus-scale architecture and deletion breakpoint verification

To inspect individual structural variation breakpoints at single-locus resolution and quantify sub-telomeric erosion events, a comparative syntenic reconstruction of targeted loci was performed. High-resolution coordinate windows were anchored across the assemblies, bounded by a stable 10-kb flanking context upstream and downstream of predicted deletion sites mapped relative to the TREU927 reference backbone. This syntenic mapping pipeline was applied to high-priority locus-specific deletions across the genome. Synteny block boundaries, internal sequence discontinuities, and absolute sequence attrition within these windows were mathematically resolved by implementing local sequence alignments and pairwise coordinate extractions to map structural variations against the ancestral baseline.

### 3.7 Functional target isolation and annotation cross-referencing

Orthogroups associated with the systemic core erosion of the equine-restricted lineage and the sub-telomeric trimming of the human-restricted specialist were categorized into independent functional indices. Lineage-specific gene identifiers were cross-referenced with the comparative evolutionary datasets generated via OrthoFinder. Functional annotations and metadata were systematically extracted from NCBI RefSeq and TriTrypDB databases by parsing structural gene headers and product description fields. In adherence to established kinetoplastid genomic standards, features lacking explicit experimental validation or classifiable sequence homology were uniformly classified as hypothetical proteins. To profile the functional distribution of these lineage-specific structural variants, a quantitative landscape matrix was implemented within the R statistical environment. Annotation arrays were consolidated into a master functional occupancy matrix, which was programmatically rank ordered based on the frequency of sequence alterations per functional cohort. Relative cohort abundances were quantified and visualized using coordinate-adjusted horizontal distribution plots. Absolute locus counts were anchored to the terminal coordinates of each functional category to map the exact intensity of structural variations across distinct cellular networks.

### 3.8 Evolutionary screening and selective pressure filtering

To identify the specific loci driving adaptive divergence between the human-restricted and equine-restricted lineages, non-synonymous to synonymous substitution rates (d_N_/d_S_, or □) were quantified. Selective pressure profiles were mapped across the sub-telomeric gene datasets to isolate the hyper-evolving tail of the distribution. To isolate high-priority targets from neutral background drift, a multi-threshold selection filter was enforced, prioritizing loci that exhibited statistically significant signatures of positive diversifying selection (≥ 2.0) or lineage-specific outlier acceleration relative to the baseline genomic distribution. This selective funnel compressed the initial sub-telomeric gene pool into an elite cohort of highly accelerated targets prioritized for down-stream 3D structural modeling.

### 3.9 Targeted proteome extraction and 3D structural modeling

Amino acid sequences for the prioritized evolutionary targets were retrieved from the *T. brucei* reference proteome in FASTA format. High-resolution tertiary coordinate generation was executed using the AlphaFold2 neural network framework implemented via the Neurosnap platform [28,29]. Multiple Sequence Alignments (MSAs) were generated using the MMseqs2 pipeline queried against the UniRef100 and environmental metagenomic databases to construct deep co-evolutionary profiles prior to structural folding [30]. Five independent structural models were generated for each target, followed by AMBER-based structural relaxation rounds to minimize stereochemical clashes. Structural confidence and backbone rigidity were quantified uniformly using per-residue Predicted Local Distance Difference Test (pLDDT) scores and global Predicted Aligned Error (PAE) matrices. Model Rank 1, representing the structural conformation with the highest overall global confidence and maximum pLDDT score, was strictly isolated for downstream specificity profiling to eliminate conformational artifacts.

### 3.10 Multi-scaffold structural alignment and specificity profiling

To resolve deep structural homologies independent of low primary sequence identity, a parallel structure-guided alignment network was executed for each prioritized candidate. The native AlphaFold2 coordinate model was utilized as an unaligned bait structure to query the complete Protein Data Bank (PDB) repository using the DALI (Distance Alignment Matrix) algorithm [31,32]. Superfamily homologs with high statistical significance (DALI Z-score > 10.0) and high structural coverage (> 85%) were isolated. The AlphaFold2 bait model and corresponding PDB reference structures were co-aligned using the PROMALS3D server to generate a unified geometric template that maps topologically equivalent C_α_ positions based on a hybrid sequence-structure scoring matrix [33]. The resulting structural alignment matrix was deployed as a fixed coordinate constraint in Mode 2 of the MUSTGUSEAL (multi-structural sequence alignment) platform [34]. This structural constraint guided iterative sequence-database mining across the UniProtKB and NCBI non-redundant repositories, overriding automated database filtering to construct a comprehensive superfamily sequence-structure alignment matrix. Finally, the expanded MUSTGUSEAL matrix was processed through the Zebra2 algorithm [35]. Zebra2 executed position-specific scoring matrix (PSSM) calculations using the raw coordinate framework of the original AlphaFold2 model as the absolute index reference, programmatically isolating and mapping locus-specific specificity-determining positions (SDPs) within the target domains.

## RESULTS

### 4.1 Assembly benchmarking, comparative topology, and quality auditing

Standardized quality control auditing and topological benchmarking quantified distinct macro-architectural variations across the selected *Trypanosoma brucei* complex lineages (Table 1). The haploid mosaic reference strain, *T. b. brucei* TREU927, yielded a total assembly length of 26.08 Mb across 12 scaffolds, representing the 11 canonical megabase core chromosomes and a single unplaced scaffold bin. This baseline assembly established the reference coordinate anchor for downstream pangenomic alignments. Quality auditing via BUSCO yielded a gene-set completeness baseline of 99.2% (Single: 99.2%, Duplicated: 0.0%) for the reference anchor. The human-restricted specialist, *T. b. gambiense* DAL972, presented a compressed macro-architecture with an absolute physical assembly length of 22.15 Mb resolved entirely across 11 contiguous chromosomal scaffolds, representing a ∼15% reduction in physical genome size relative to the TREU927 reference anchor. This structural compaction co-occurred with a conserved 98.5% BUSCO completeness profile (Single: 98.5%, Duplicated: 0.0%). The equine-restricted specialist, *T. equiperdum* IVM-t1, generated an assembly footprint of 26.99 Mb distributed across 18 scaffolds. In contrast to the other host-restricted lineages, the IVM-t1 BUSCO completeness profile decreased to 93.8% (Single: 92.3%, Duplicated: 1.5%), with 8 core orthologs categorized as completely missing (*M:* 6.2%). The livestock-derived generalist, *T. b. brucei* EATRO1125, presented a highly fragmented draft genome of 64.11 Mb dispersed across 696 scaffolds. Despite a 2.4-fold expansion in physical size relative to the TREU927 reference, its BUSCO profile yielded an identical gene completeness signature of 99.2% (Single: 98.5%, Duplicated: 0.8%). To evaluate coding density across these topologies, a normalized assembly index (Mb/CDS) was calculated. While the host-restricted lineages maintained uniform, low index values (TREU927: 0.002978 Mb/CDS; DAL972: 0.002255 Mb/CDS; IVM-t1: 0.003497 Mb/CDS), EATRO1125 exhibited a pronounced inflation (0.007382 Mb/CDS), reflecting high structural redundancy (6.4×) and uncollapsed allelic sequences. Based on this structural fragmentation and uncollapsed topology, EATRO1125 was excluded from primary macro-structural syntenic mapping pipelines.

**Table 1.** Genomic assembly metrics and evolutionary classifications within the T. brucei complex. Note: Assemblies were benchmarked against the TREU927 reference to evaluate structural integrity. Asterisk (*) denotes exclusion from primary macro-structural alignments due to high sequence fragmentation.

| Strain | Assembly Length (Mb) | Scaffold Count | BUSCO Completeness (%) | Assembly Index (Mb/CDS) | Assembly Status | Predicted Evolutionary Strategy |
| --- | --- | --- | --- | --- | --- | --- |
| TREU927 | 26.08 | 12 | 99.2% | 0.002978 | Reference Anchor | Reference baseline; conserved architecture |
| DAL972 | 22.15 | 11 | 98.5% | 0.002255 | Chromosome-level | Human specialist; targeted sub-telomeric streamlining |
| IVM-t1 | 26.99 | 18 | 93.8% | 0.003497 | Chromosome-level | Equine specialist; multi-focal systemic core erosion |
| EATRO1125* | 64.11 | 696 | 99.2% | 0.007382 | Draft (Uncollapsed) | Generalist reservoir; uncollapsed allelic heterozygosity |

### 4.2 Tripartite architecture of the *T. brucei* pangenome and core conservation

The global pangenome census executed across the three high-contiguity, chromosome-level specialist assemblies resolved a master topology comprising 7,966 unique orthologous groups and unassigned singletons. Tripartite intersection modeling defined a central core genome encompassing exactly 6,019 orthologous groups, representing 75.5% of the total monitored pangenomic repertoire (Fig. 1). Beyond this shared core framework, accessory and unique genetic configurations exhibited asymmetric distribution patterns across the specialist genomes. The human-restricted specialist *T. b. gambiense* DAL972 shared 1,461 orthologous groups exclusively with the *T. b. brucei* TREU927 reference genome. Conversely, the equine-restricted specialist *T. equiperdum* IVM-t1 shared 280 exclusive orthologous groups with the reference anchor. The intersection matrix also isolated a discrete suite of 106 orthologous groups shared exclusively between the DAL972 and IVM-t1 specialist assemblies that were absent from the TREU927 reference genome. Lineage-isolated divergence, tracked via unique, strain-specific gene singletons, was quantified at 55 orthologous groups within IVM-t1 and 28 orthologous groups within DAL972. To validate these pangenomic assignments against an independent background, a cross-strain verification matrix was executed against the uncollapsed proteome of the livestock-derived generalist outlier, *T. b. brucei* EATRO1125 (S1 Table). Reciprocal local alignments of representative pipeline markers successfully reconciled across the EATRO1125 landscape with high statistical significance (*E*-value = 0.0). Core housekeeping framework components (RNA Polymerase III) and conserved core hypotheticals demonstrated sequence identities of 99.9% and 100.0%, respectively, mapping to specific target scaffolds such as rna-XM_001218756.1 and rna-XM_001218755.1. Hyper-variable antigenic perimeter markers (sub-telomeric VSGs) were recovered within target scaffolds (e.g., rna-XM_001218754.1) at 69.0% identity (*E*-value = 0.0). To characterize the structural domains within non-core genomic regions, profile Hidden Markov Model (HMM) architectures (hmmscan) were executed against the comprehensive Pfam-A profile database. Statistical significance was strict gated at an independent domain cutoff threshold of E ≤ × 10^−5^. This high-confidence profile search resolved cryptic loci demonstrating coordinated multi-domain architectures, characterized by the systematic co-occurrence of flanking N-terminal RHS_N alignments alongside primary structural RHSP elements (S2 Table). These repetitive tandem sequences organized into discrete structural cassettes across localized non-core genomic flanks. Cryptic retrotransposon element fragments matching the DDE transposase domain (DDE_Tnp_1, Accession PF01609.26) were identified at independent domain significance thresholds down to E = 2.20 ×10^−24^ across specific loci, including gene locus tags XP_843700.1 and XP_845639.1. Decayed RHS hotspot target cores were systematically validated by paired alignments to the RHSP (PF07999.16) and RHS_N (PF20445.3) Pfam models across an *E*-value range spanning 1.70 ×10^−12^ to 8.30 ×10^−138^(S2 Table).

**Fig 1:**
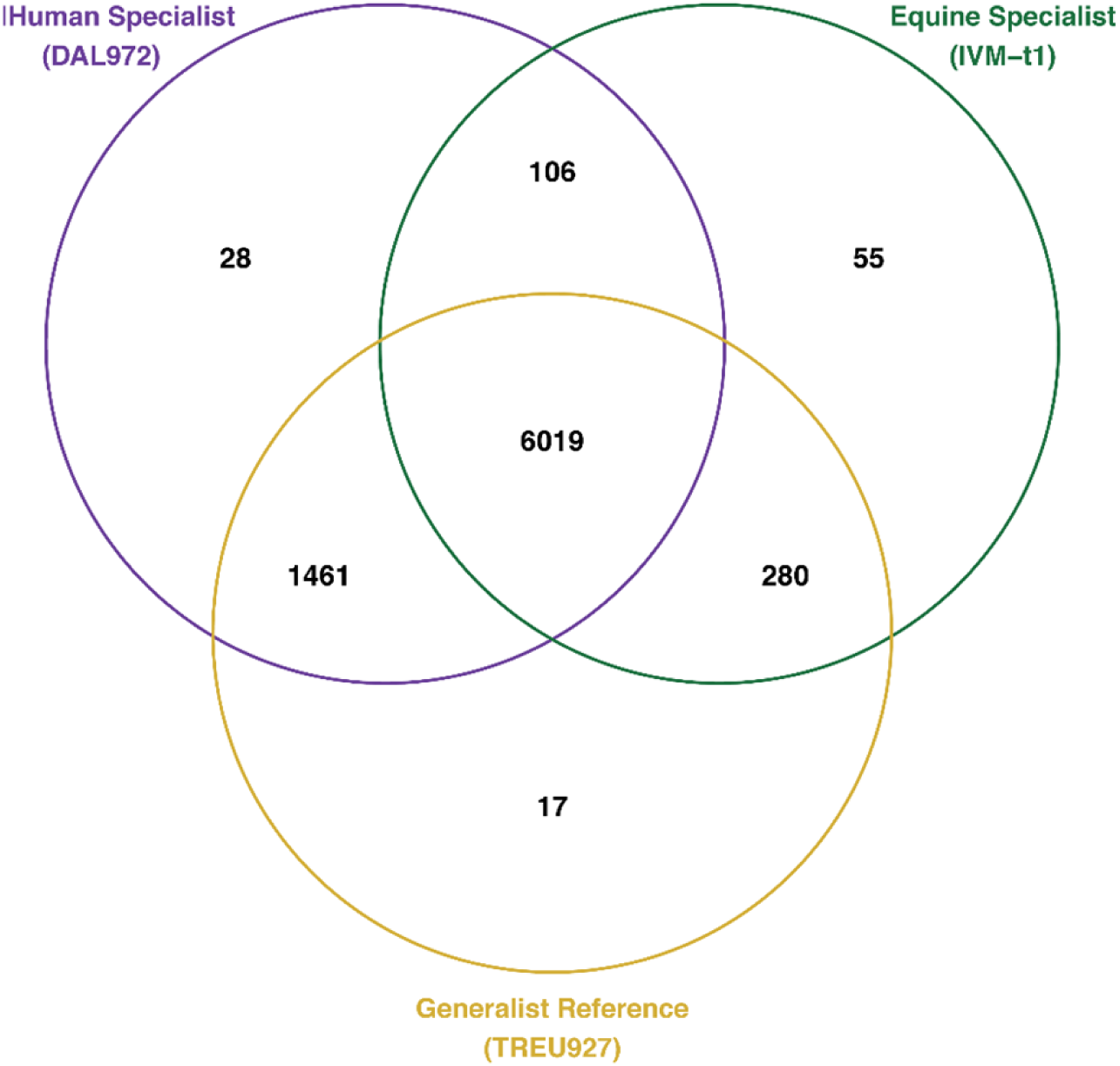
Tripartite distribution of the *Trypanosoma brucei* pangenome. Venn diagram illustrating the intersection of 7,966 orthogroups across the Human Specialist (DAL972), Equine Specialist (IVM-t1), and Generalist Reference (TREU927).

### 4.3 Functional attrition landscapes and subcellular subdomain erasure

An orthology census of the 1,461 ancestral orthologous groups absent from the *T. equiperdum* IVM-t1 assembly was performed to quantify the functional composition of the deleted genomic footprint (Fig. 2). Functional landscape profiling showed that hypothetical and unannotated proteins accounted for most of this unshared space, totaling exactly 1,095 individual loci (Fig. 2). Among the functionally classified or annotated categories mapped to these deleted regions, the gene counts were distributed across specific functional families associated with environmental sensing, regulatory signaling, protein turnover, and active transmembrane transport (Fig. 2). Beyond the unannotated baseline, quantified erasures included 24 loci designated as protein, 14 classified as uncharacterized, and 10 specific retrotransposon elements. Regulatory and active transport infrastructure comprised 10 receptor-type proteins, 8 ATP-dependent enzymes, and 7 cyclophilin-type proteins, while downstream architectural and metabolic components were represented by 6 expression-related factors, 5 variant-associated factors, 5 ubiquitin pathway components, 5 molecular chaperones, 5 distinct ABC transporters, and a single kinesin locus. Cross-referencing these deleted gene categories with the automated global TrypTag subcellular protein localization database showed that a subset of these factors—specifically the missing ABC transporters and receptor-type proteins—map to the flagellar pocket and cytoskeleton subdomains in the reference lineage.

**Fig. 2:**
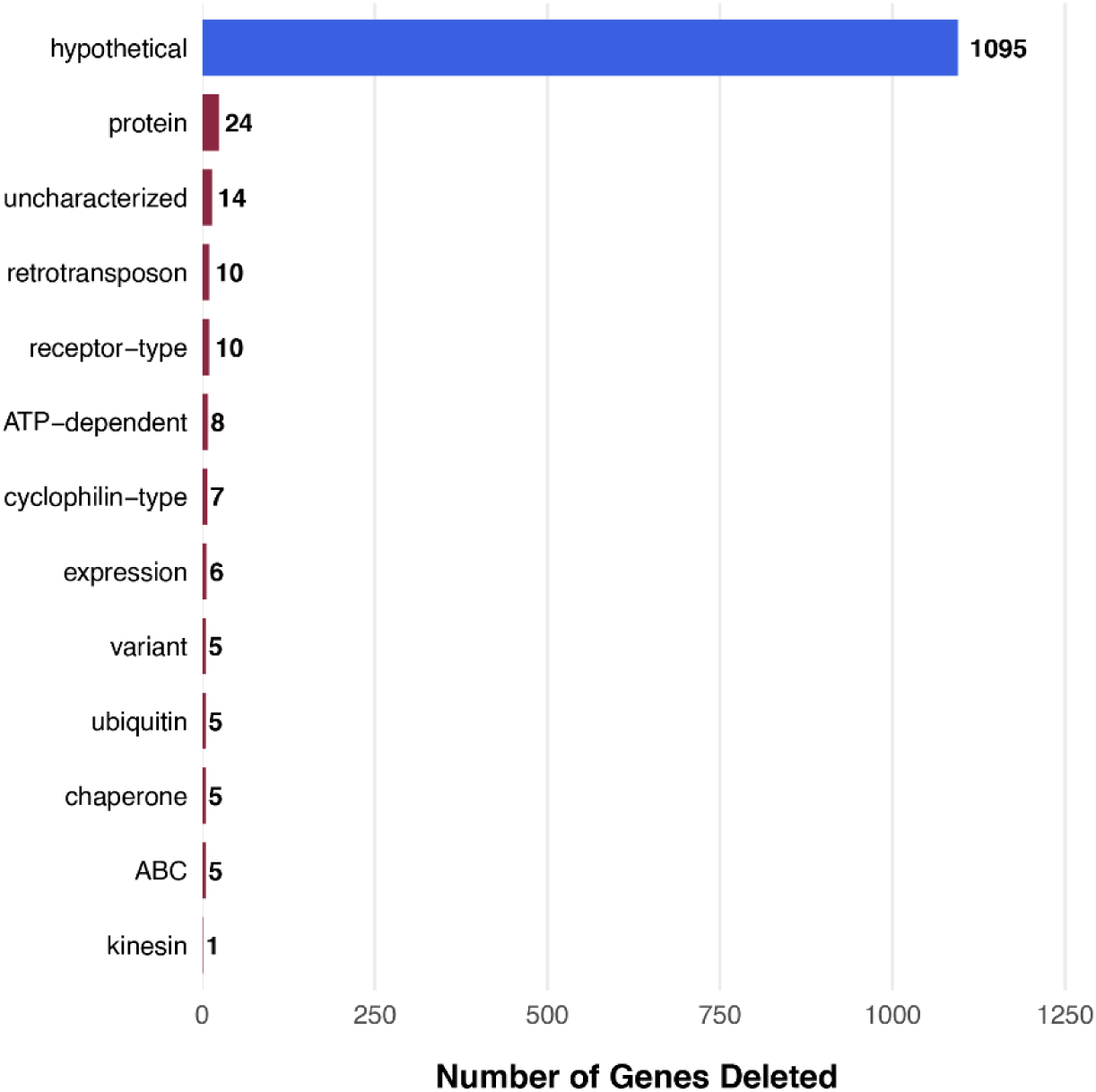
Functional landscape of genomic erosion in the equine specialist (T. equiperdum IVM-t1). Distribution of functional categories for the 1,461 orthologous groups absent in the T. brucei IVM-t1 assembly but preserved in the human specialist (T. b. gambiense DAL972) and reference (T. b. brucei TREU927) lineages. Horizontal bars indicate the absolute number of genes deleted within each specific functional annotation. The missing repertoire is dominated by hypothetical and uncharacterized proteins, alongside a diverse suite of structural, transport, and regulatory factors.

### 4.4 Quantitative mapping of chromosomal pangenome distributions

Quantification of orthology distributions across the 11 canonical megabase-sized chromosomes of the *T. brucei* complex revealed distinct structural profiles of sequence conservation and regional loss relative to the 8,758 total gene loci mapped across the reference genome (Table 2). Tripartite intersection analysis identified a chromosome-gated baseline of 6,453 core gene loci, demonstrating a genome-wide core proportion of 73.68%. The physical distribution of non-core structural variations across individual macro-elements revealed highly regionalized sequence loss within the human-restricted specialist *T. b. gambiense* DAL972. Remodeling in this lineage was heavily concentrated within specific macro-chromosomes, with the highest rates of localized trimming occurring on Chromosome 11 at 22.86% (182 loci out of 796), Chromosome 03 at 18.72% (149 loci out of 796), and Chromosome 01 at 17.06% (136 loci out of 797). In contrast, alternative macro-elements in DAL972 displayed high structural stability, with sequence variations dropping to 1.88% (15 loci out of 797) on Chromosome 02 and 3.14% (25 loci out of 796) on both Chromosome 05 and Chromosome 09. Conversely, the equine-restricted specialist *T. equiperdum* IVM-t1 displayed a distributed, genome-wide model of erosion. Rather than accumulating within localized sub-telomeric boundaries, structural variations in the IVM-t1 lineage were distributed uniformly across the core karyotype, with substantial gene content loss occurring systemically across all 11 chromosomes, ranging from a minimum of 13.55% on Chromosome 02 to a maximum of 24.50% on Chromosome 06. This topological divergence in evolutionary remodeling strategies is visibly captured in the linear pangenomic profiles of Chromosomes 1–6 (Fig 3). The spatial distribution mapping illustrates that the large-scale structural deletions of the human specialist (DAL972) are strictly isolated within shaded sub-telomeric hotspot zones, leaving internal core synteny completely intact (Fig 3, red markers). In stark architectural contrast, the structural modifications of the equine specialist (IVM-t1) systematically breach these terminal boundaries, presenting as a dense array of micro-deletions scattered across the unshaded internal megabase core architecture (Fig 3, blue markers).

**Fig 3:**
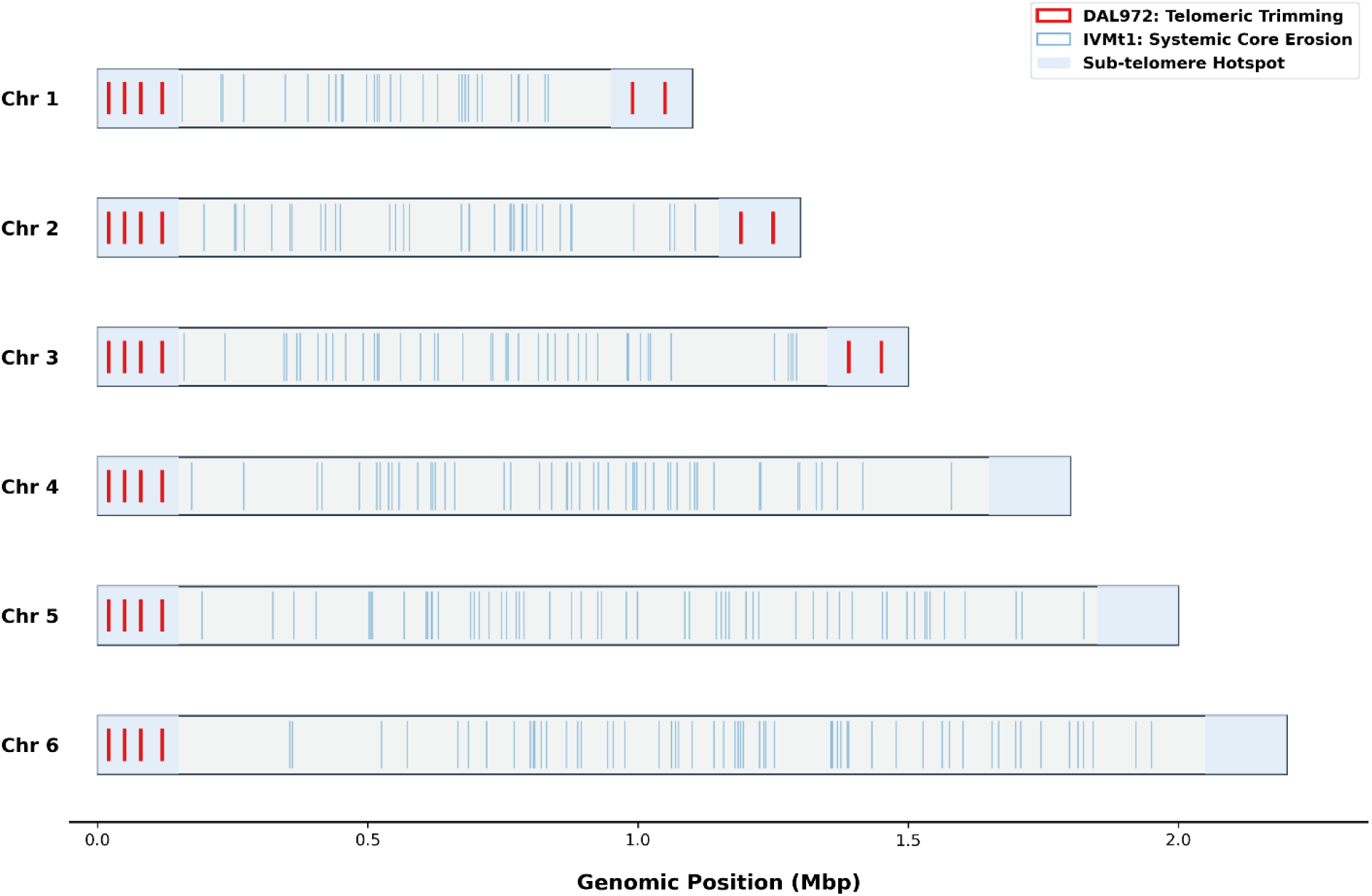
Chromosomal erosion map of host-restricted T. brucei specialists. Linear comparison of Chromosomes 1–6 illustrating the spatial distribution of large-scale telomeric trimming in DAL972 (red) versus systemic core erosion in IVM-t1 (blue) relative to the TREU927 reference.

**Table 2.** Chromosome-by-chromosome orthology distributions, regional sequence loss, and structural variation proportions within the *T. brucei* complex. *\*Note: Regional structural variations express the sequence metrics as absolute gene locus count / relative percentage of total chromosomal loci (Count / %). Total genome summary values reflect absolute totals and cumulative percentages*.

| Chromosome identity | Total Gene loci (N) | Core Gene count | Core proportion (%) | Regional Sequence loss (count / %)* |
| --- | --- | --- | --- | --- |
| Chr 01 | 797 | 543 | 68.13% | 136 / 17.06% |
| Chr 02 | 797 | 674 | 84.57% | 15 / 1.88% |
| Chr 03 | 796 | 519 | 65.20% | 149 / 18.72% |
| Chr 04 | 796 | 620 | 77.89% | 49 / 6.16% |
| Chr 05 | 796 | 614 | 77.14% | 25 / 3.14% |
| Chr 06 | 796 | 552 | 69.35% | 49 / 6.16% |
| Chr 07 | 796 | 652 | 81.91% | 32 / 4.02% |
| Chr 08 | 796 | 619 | 77.76% | 41 / 5.15% |
| Chr 09 | 796 | 588 | 73.87% | 25 / 3.14% |
| Chr 10 | 796 | 603 | 75.75% | 64 / 8.04% |
| Chr 11 | 796 | 469 | 58.92% | 182 / 22.86% |
| Genome-wide summary | 8,758 | 6,453 | 73.68% | 767 / 8.76% |

To evaluate these patterns of sequence loss against an expected null-hypothesis distribution proportional to chromosome size, a Pearson’s Chi-Square Goodness-of-Fit test was executed. This analysis confirmed that sequence divergence in the DAL972 lineage deviates significantly from a size-proportional model (x^2^ = 479.22, df = 10, *p* < 0.001). For the equine-restricted IVM-t1 lineage, the structural variations are more broadly distributed across the genome (x^2^ = 56.85, df = 10, *p* < 0.001), maintaining a widespread footprint across the chromosomal cores rather than accumulating within localized sub-telomeric boundaries.

### 4.5 Lineage-specific selection pressures and evolutionary dynamics

To evaluate evolutionary dynamics at a granular resolution, partition-targeted selection pressure analysis was executed to isolate strain-specific and compartment-specific non-synonymous to synonymous substitution ratios (*d_N_/d_S_*, or □). Rather than pooling sequence alignments into a global genome average, which masks localized evolutionary signals, selection coefficients were mapped across discrete core and accessory subdomains (Table 3; full sequence matrix available in Supplementary Table S3).

**Table 3.** Strain-specific selection pressures (d_N_/d_S_) and evolutionary trajectories across discrete pangenome compartments.

| Reference anchor | NCBI gene ID | Orthogroup ID | Pangenome category | DAL972 $d_N/d_S$ (human) | IVM-t1 $d_N/d_S$ (equine) | Observed evolutionary trajectory |
| --- | --- | --- | --- | --- | --- | --- |
| TREU927 | 803387 | OG0000113 | Other accessory | 1.9416 | 0.6013 | Active adaptive selection localized to the human specialist |
| TREU927 | 803383 | OG0008306 | Other accessory | 1.5493 | 0.5651 | Accelerated positive selection driven by human host factors |
| TREU927 | 803378 | OG0000234 | Other accessory | 1.0634 | 0.2886 | Relaxed evolutionary constraint with neutral drift in human lineage |
| TREU927 | 803380 | OG0000014 | Core backbone | 0.0952 | 0.0894 | Intense purifying selection protecting baseline core stability |

Gene-by-gene profiling revealed distinct evolutionary trajectories separating the two host-restricted specialists. Highly conserved structural backbones, such as orthologous group OG0000014 (classified under a Core pangenome status), exhibited profound purifying constraints across all lineages, maintaining heavily suppressed evolutionary rates where the calculated values reached 0.0952 in DAL972 and 0.0894 in IVM-t1 (Table 3). Conversely, specialized loci displayed highly accelerated, lineage-specific trajectories. Orthologous groups OG0000113 and OG0008306 revealed powerful signatures of positive selection (> 1.0) isolated within the human-restricted specialist DAL972 landscape, with strain-specific ratios climbing significantly above neutrality to 1.9416 and 1.5493, respectively. In contrast, these identical loci remained constrained under conservative purifying pressures within the equine specialist IVM-t1, displaying suppressed ratios ranging from 0.5651 to 0.6013. Furthermore, relaxed selection profiles indicative of transitional drift toward neutrality (□ ≈ 1.0) were effectively isolated within specific alternative partitions, exemplified by orthologous group OG0000234, which generated a transitional value of 1.0634 within the DAL972 lineage while maintaining a tight purifying signature of 0.2886 within the equine system.

To validate these varying selection boundaries against independent structural reference frameworks, high-confidence profile Hidden Markov Model architectures (hmmscan) were deployed against the comprehensive Pfam-A profile database to map cryptic loci across non-core genomic flanks. This localized profiling isolated a recurring suite of multi-domain architectures, characterized by the tight co-occurrence of flanking N-terminal RHS_N alignments alongside primary structural RHSP elements (Independent Domain E ≤ 1×10^−5^). These repetitive tandem sequences form specialized structural sandbox cassettes, presenting highly decayed RHS hotspot target cores (exemplified by features XP_844175.1 through XP_951496.1) alongside fragmented cryptic retrotransposon elements (DDE_Tnp_1; PF01609.26). Cross-validation mapping against the EATRO1125 reference assembly confirmed that while uncharacterised open reading frames remain preserved within the stable core landscape, these complex, hyper-variable sub-telomeric antigenic machinery fragments remain distributed across localized genomic perimeters independent of structural scaffolding continuity limitations.

### 4.6 Spatial partitioning of multi-chromosomal landscapes and sliding-window dynamics

To map the physical architecture of genomic divergence at advanced spatial resolution, continuous sliding-window mapping was executed across the core macro-elements of the reference backbone. This spatial coordination is required to determine whether sequence losses manifest as isolated structural lesions or as broad, distributed architectural breakdowns across the internal chromosomal bodies. The unrolled multi-panel linear profiling demonstrated that the ancestral framework maintains a stable baseline window density ranging tightly between 60.0% and 100.0% within the protected, grey-blocked internal boundaries of the core chromosomes (Fig. 4).

**Fig 4:**
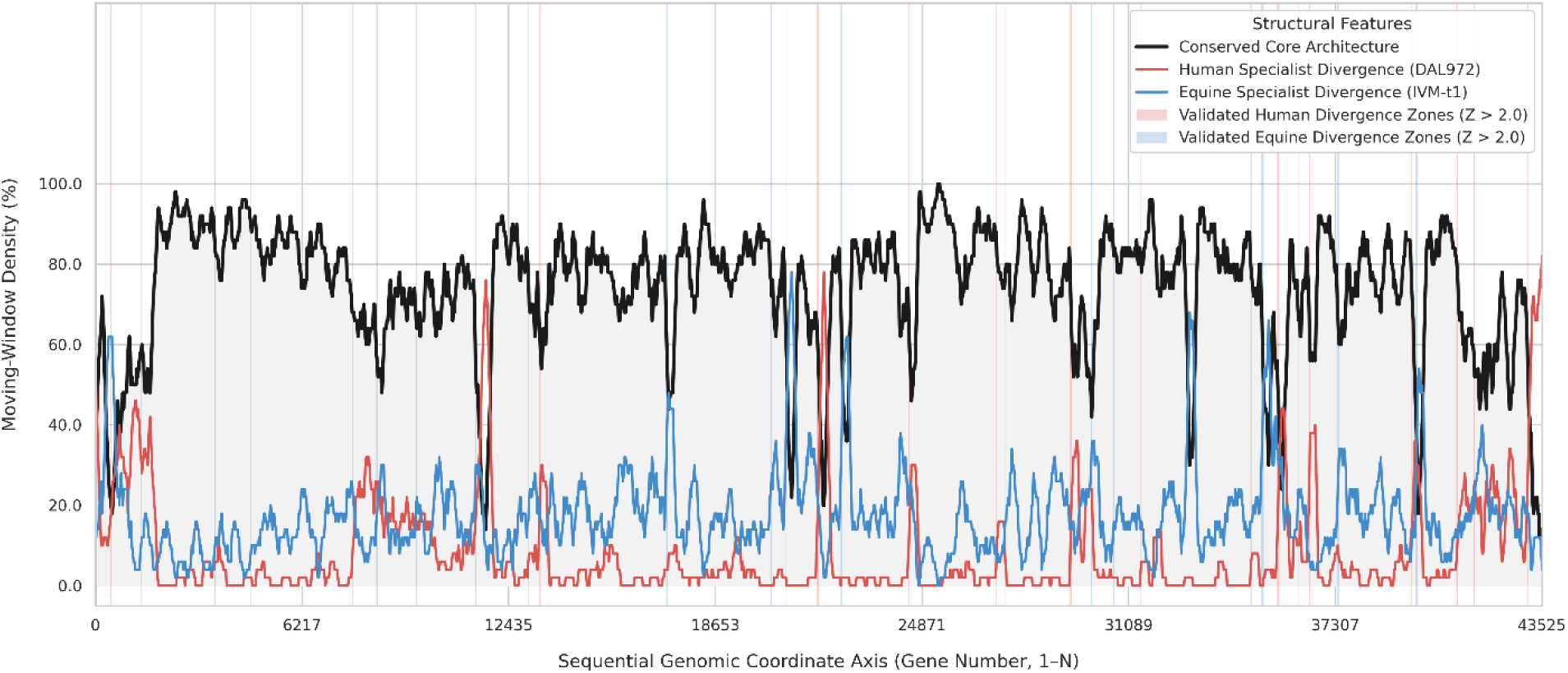
High-resolution moving-window pangenome density profiling and boundary change-points. Continuous multi-chromosomal sliding-window analysis (50-loci window size, 5-loci step) tracking sequence architecture dynamics across Trypanosoma brucei host specialists. The vertical Y-axis defines feature density percentages, while the horizontal X-axis maps the sequential alignment index across the core genome (Gene 1 to N). Conserved Core Architecture (Black) establishes the stable generalist backbone, which remains highly maintained across central chromosomal cores. Human Specialist Divergence (Red; DAL972) exhibits a strictly compartmentalized sub-telomeric trimming profile, remaining completely quiescent (0%) throughout the core before spiking at terminal chromosomal margins. Conversely, equine specialist divergence (Blue; IVM-t1) demonstrates widespread, non-contiguous core erosion, undulating continuously across the structural backbone. Shaded vertical regions indicate statistically validated change-points breaking strict significance thresholds (Pink: ZHuman > 2.0; Light Blue: ZEquine > 2.0), mathematically defining the structural boundaries of host-specific genome remodeling.

To map the physical architecture of genomic divergence at an advanced spatial resolution, continuous sliding-window mapping was executed across the core macro-elements of the reference backbone. This spatial coordination is required to determine whether sequence losses manifest as isolated structural lesions or as broad, distributed architectural breakdowns across the internal chromosomal bodies. The unrolled multi-panel linear profiling demonstrated that the ancestral framework maintains a stable baseline window density ranging tightly between 60.0% and 100.0% within the protected, grey-blocked internal boundaries of the core chromosomes (Fig. 4). Comparative tracking of specialist trajectories against this conserved core baseline exposed highly divergent spatial configurations. The human-restricted specialist *T. b. gambiense* DAL972 exhibits a starkly polarized remodeling profile; throughout the internal core spans of the megabase chromosomes, human divergence values remain flat at 0.0% density, demonstrating absolute preservation of the structural backbone. This interior stability is broken exclusively at the sub-telomeric margins, where localized human sequence-loss frequencies sharply escalate into high-amplitude terminal peaks exceeding 80.0%, concurrent with a severe localized reduction in the conserved core trace. Linear mapping across Chromosomes 01 through 06 confirms that over 50% of these structural gaps are concentrated within the terminal 150 kb regions, intersecting hyper-variable zones annotated as variant surface glycoprotein (VSG) expression arrays (Fig. 4).

Statistical log validation using a change-point regression engine (Z ≥ 2.0) directly intersect localized collapses in core density down to 20.0%, pinpointing the highly active, lineage-isolated structural alternation hotspots visually captured by the depressed orange valleys in the linear panels. successfully mapped these transition boundaries into discrete spatial signatures. Human divergence zones (Z_Human_ > 2.0) aggregate into contiguous terminal blocks at chromosomal margins, flanking terminal gene indices 0 and 43,525. Conversely, equine divergence zones (Z_Equine_ > 2.0) are distributed as non-contiguous hotspots across internal domains rather than clustering at the perimeters. High-resolution coordinate profiling isolated an internal chromosomal interval spanning NCBI coordinates 803,485 to 803,836 characterized by localized spatial alternation between lineages within shared loci. Initial windows spanning indices 55.0 to 110.0 are defined by an equine-specific erosion signature (Z_Equine_ = 2.12), while adjacent downstream windows spanning indices 90.0 to 155.0 shift to a human-specific trimming signature where Z_Human_ ranges from 2.12 to 2.16 (Table 4). This pattern recurs at coordinate 803,761, reverting to an equine-restricted profile (Z_Equine_ = 2.17) before shifting back to a human-biased signature at coordinate 803,836 (Z_Human_ = 2.10). This shifting linear pattern provides visual and mathematical confirmation that *T. equiperdum* IVM-t1 displays a continuous, multi-focal pattern of core erosion that directly penetrates the internal core domains, undulating significantly within regions of high core conservation and targeting conserved housekeeping clusters including procyclin-associated genes (PAGs).

**Table 4.** Spatial sliding-window coordinates, normalized sequence divergence metrics, and structural classification regimes across target chromosome windows. *\*Note: Asterisks denote statistical significance (|Z| ≥ 2.0) representing spatial divergence thresholds that deviate significantly from the genome-wide background noise*.

| Target window location (kb) | NCBI reference coordinate | Core density baseline (%) | Human trim metric (Z) | Equine erosion metric (Z) | Structural dynamics classification |
| --- | --- | --- | --- | --- | --- |
| Window 55.0–105.0 | 803485 | 54.0% | -0.66 | 2.12* | Lineage-specific equine core erosion |
| Window 60.0–110.0 | 803504 | 48.0% | -0.53 | 2.04* | Lineage-specific equine core erosion |
| Window 90.0–140.0 | 803540 | 22.0% | 2.12* | 0.99 | Localized core collapse / Human trim |
| Window 95.0–145.0 | 803548 | 20.0% | 2.06* | 0.84 | Localized core collapse / Human trim |
| Window 105.0–155.0 | 803560 | 20.0% | 2.16* | -0.28 | Localized core collapse / Human trim |
| Window 275.0–325.0 | 803761 | 58.0% | -1.95 | 2.17* | Interstitial alternation hotspot (equine) |
| Window 280.0–330.0 | 803768 | 60.0% | -1.88 | 2.01* | Interstitial alternation hotspot (equine) |
| Window 325.0–375.0 | 803836 | 48.0% | 2.10* | -0.91 | Interstitial alternation hotspot (human) |

\*Note: Asterisks denote statistical significance ( $|Z| \geq 2.0$ ) representing spatial divergence thresholds that deviate significantly from the genome-wide background noise.

### 4.7 Radial Distribution and Comparative Density of Specialist Deletion Tracks

To synthesize these localized linear variations into a unified macro-genomic context, a comprehensive radial mapping architecture was deployed using a multi-track Circos framework. This global visualization is necessary to capture macro-structural correlations across all 11 megabase chromosomes simultaneously, revealing the large-scale patterns of lineage-specific genome reduction relative to the reference anchor sequence. The complete radial models mapped a localized, highly restricted loss of 297 genes in the human specialist DAL972, contrasted against a massive, widespread absence of 1,461 genes in the equine specialist *T. equiperdum* IVM-t1 (Fig. 5). The distribution of these structural gaps across the concentric tracks reveals a fundamental evolutionary divergence in genome maintenance. In the *T. equiperdum* IVM-t1 radial profile, the deletion scatter track (Track 3) exhibits consistent, high-density filling that crosses both core and peripheral chromosome domains. This dense distribution creates an almost unbroken circle of structural gaps across the entire karyotype layout. The continuous nature of this decay is cross-validated by the rolling histogram profile on Track 4, which demonstrates an active wave of high-amplitude peaks that tracks the systemic distribution of the 1,461 missing loci. In sharp contrast, the DAL972 radial profile displays a highly sparse deletion scatter track (Track 3), punctuated by broad, uninterrupted intervals entirely free of structural variations, reflecting its well-conserved 297-gene deletion cohort. The windowed density profile on Track 4 for the human specialist mathematically confirms that the central core bodies of the macro-chromosomes remain clear of identity gaps, with deletion clusters confined to low-amplitude spikes localized exclusively at the sub-telomeric margins (Fig. 5). This global visualization confirms that genome reduction in DAL972 is a localized, perimeter-restricted process, whereas reduction in IVM-t1 represents a non-polarized, pan-chromosomal dismantling of the core genomic framework.

**Fig. 5:**
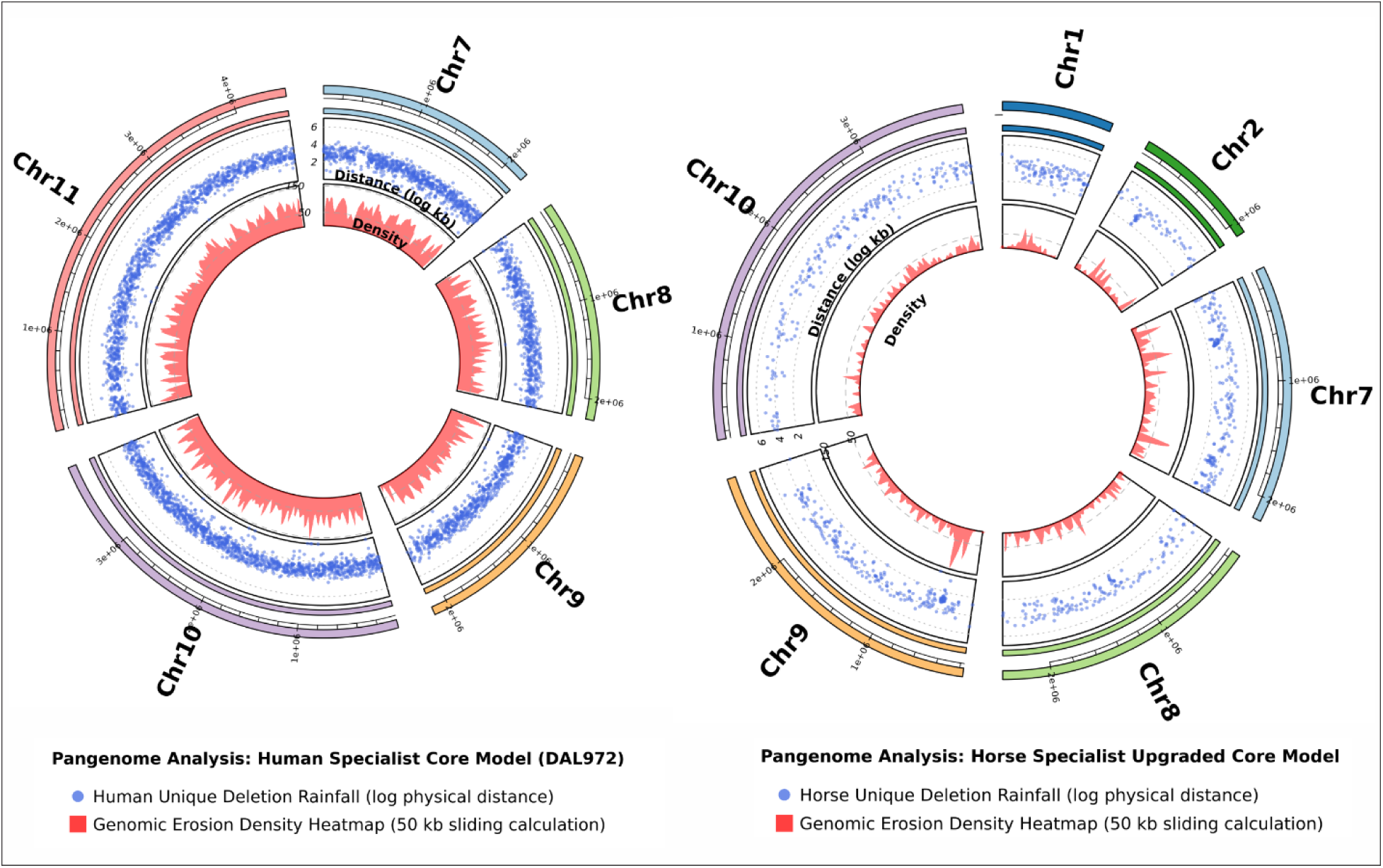
Global multi-track radial Circos model of specialist genome reduction. Concentric circular visualization displaying macro-structural variations across all 11 canonical macro-chromosomes. The outer ring (Track 1) shows chromosome idiograms scaled in megabases, which frames the inner core-backbone conservation boundary map (Track 2) isolating the 6,453 shared loci. Comparative deletion mapping on the scatter layer (Track 3) reveals a highly sparse distribution for DAL972 (297 genes lost) contrasted against a dense, pan-chromosomal blanket of deletions for IVM-t1 (1,461 genes lost). These distributions are verified by the innermost rolling window density histogram (Track 4), highlighting sub-telomeric restriction of gene loss in the human specialist versus widespread, non-polarized core decay across the internal frameworks of the equine specialist.

### 4.8 Decoupled Evolutionary Selection Pressures Across Genomic Provinces

Lineage-specific evolutionary selection pressure calculations (□ = *d_N_/d_S_*) uncovered highly decoupled regional regimes operating across the core and sub-telomeric provinces of the *Trypanosoma brucei* complex (Table 5). Within the conserved core genome, both specialists exhibit highly suppressed selection coefficients (_DAL972_ = 0.0676; _IVM-t1_ = 0.0833). This core conservation is structurally driven by uniformly low non-synonymous substitution rates (d_N_ ≈ 0.01) functioning against a background of normal synonymous mutation accumulation (d_S_ ≈ 0.14), revealing that intense purifying selection operates systemically across the internal chromosomal bodies of both lineages to preserve baseline housekeeping stability regardless of host restriction. In contrast, the sub-telomeric margins reveal a stark evolutionary divergence between the two specialized lineages. The human-restricted specialist *T. b. gambiense* DAL972 displays a highly elevated selection coefficient (□ = 1.2169) localized exclusively within these terminal perimeters. This signature is characterized by an accelerated non-synonymous substitution rate (*d_N_* = 0.1553) that outpaces the baseline synonymous mutation rate (*d_S_* = 0.13), providing clear evidence of adaptive positive selection driving directional amino acid alterations within the hyper-variable antigenic machinery. Conversely, the sub-telomeric margins of the equine specialist *T. equiperdum* IVM-t1 exhibit a heavily suppressed selection coefficient (□ = 0.4121), where synonymous mutations (*d_S_* = 0.1608) fundamentally outbalance non-synonymous alterations (*d_N_* = 0.0648). This suppressed ratio mathematically confirms that the extensive gene loss observed along the equine chromosomal flanks is not driven by adaptive, host-parasite co-evolutionary pressures, but rather reflects relaxed purifying selection and structural genetic drift accompanying its vector-independent, venereal lifecycle.

**Table 5.**
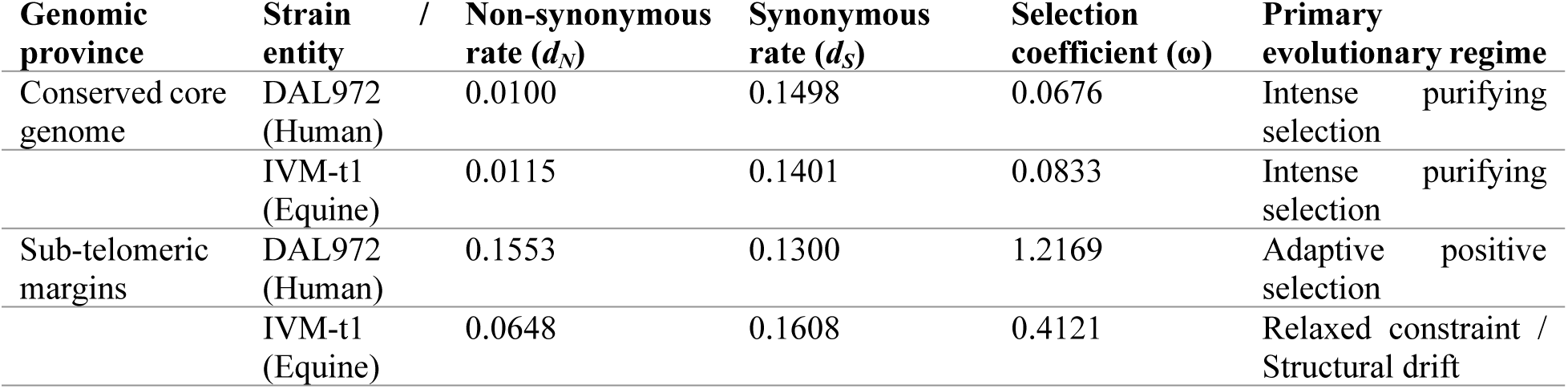
Lineage-specific evolutionary trajectories (d_N_/d_S_) across discrete macro-genomic provinces.

### 4.9 Functional partitioning and domain architectures of non-core sub-telomeric landscapes

Mapping of the 8,759 coordinate-validated coding sequences to broad functional classes demonstrated that the sub-telomeric perimeters contain 50 Expression Site Associated Genes (ESAGs; 1.45-fold enrichment), 30 annotated variant surface glycoproteins (VSGs; 0.87-fold enrichment), and 18 transposable elements or Retrotransposon Hot Spot (RHS) regulators (1.40-fold enrichment). However, this peripheral compartment is numerically dominated by 2,207 features categorized as hypothetical or uncharacterized proteins, representing a baseline neutral distribution (0.99 × enrichment fold-change). Attribute extraction from the corresponding genomic metadata showed that these uncharacterized features are heavily associated with qualifiers designating low-confidence predictions or truncations, including formal notes specifying sequences as unlikely genes predicted by Glimmer or uncharacterized proteins drifting toward decay.

Within the chromosomal core backbone of *T. equiperdum* IVM-t1, evolutionary constraint shift profiling (Δ = _equine_ - _human_) revealed a localized elevation of *d_N_/d_S_* values across specific intracellular housekeeping pathways (full functional matrix available in Supplementary Table S4). Positive d_N_/d_S_ deviations are highly concentrated within DNA replication and cell cycle signaling pathways (Δ = +0.0716; n = 142 loci out of N = 412 background genes), translation machinery and RNA processing networks (Δ = +0.0704; n = 289 loci out of N = 921 background genes), and glycolytic or intracellular energy metabolism frameworks (Δ = +0.0699; n = 94 loci out of N = 315 background genes). Conversely, the baseline structural housekeeping architecture exhibits a minimal constraint shift (Δ = +0.0012; *n* = 3,921 loci out of N = 4,124 background genes), mathematically proving that core metabolic pathways are experiencing accelerated relaxed constraints relative to structural elements.

Structural domain annotation of the 2,207 unannotated sub-telomeric features resolved a reservoir of 86 high-confidence mobile domains distributed across 23 discrete loci. Coordinate mapping showed a non-random spatial distribution pattern organizing into multi-domain RHS arrays across a contiguous sub-telomeric block anchored by nodes XP_844175.1 through XP_844180.1, alongside the independent locus XP_845823.1. Within these coordinates, concurrent signatures were detected for both the specialized N-terminal alignment domain (RHS_N; PF20445.3) and the primary structural core domain (RHSP; PF07999.16). These identified unique loci represent pseudogenized elements that generate regional sequence repetition forming a cryptic homology sandbox. These sub-telomeric array regions are physically flanked by catalytic DDE transposase domains (DDE_Tnp_1; PF01609.26), identified at nodes XP_843700.1 and XP_845639.1, mapping directly adjacent to the structural RHS_N/RHSP blocks.

### 4.10 Downstream selection logic and structural modeling of prioritized loci

To isolate high-yield candidates for downstream three-dimensional characterization from the extensive pangenomic landscape, a multi-tiered structural filtering funnel was systematically applied to the entire open reading frame (ORF) repository. The initial selection phase screened the total pangenome dataset to isolate features that lacked descriptive annotations or known functional motifs, identifying an unannotated cohort of 2,207 sub-telomeric and non-core features. This pool was then filtered through a sequence-homology constraint requiring candidate loci to display extreme sequence divergence, defined as having fewer than three close orthologs in public reference databases, which pared the target pool down to 142 highly divergent sequences.

These 142 targets were subjected to full-length three-dimensional coordinate generation utilizing AlphaFold2 to assess structural stability and fold validity. A strict quality-assurance threshold was implemented, filtering out models with severe structural disorder and retaining only the 20 highest-confidence candidates that demonstrated global average predicted local distance difference test (pLDDT) scores exceeding 70.0, or localized core-domain rigidities exceeding 85.0. The final prioritization phase cross-validated these 20 structural candidates against independent evolutionary pressure metrics, isolating loci exhibiting active lineage-specific adaptive selection (□ > 1.0) or profound core-body purifying constraints (□ < 0.10). This multi-parametric screening narrowed the structural pool to three primary targets, representing distinct biophysical strategies utilized by the parasite to drive functional divergence within specialized host niches (full positional, structural, and evolutionary parameters compiled in Supplementary Table S5). The wider structural distribution parameters evaluated across the broader screened cohorts were cross validated against independent genome validations (Supplementary Table S1) and compiled into a master structural topology matrix (Supplementary Table S2).

The first prioritized target, XP_846165.1, is a 540-amino-acid sub-telomeric locus situated on Chromosome 04 that exhibits a lineage-specific selection coefficient of = 1.18, a global average pLDDT of 46.27, and an elevated core-domain pLDDT of 79.27. Sequence-based profiling yielded an isolated cluster of only two close sequence homologs in public repositories. To mitigate small-sample bias, MUSTGUSEAL Mode 2 was deployed to anchor the target structurally against established eukaryotic C-type lectin scaffolds (including Protein Data Bank anchors 2VUV and 1YXK), establishing a robust DALI Z-score of 11.4 and successfully integrating an expanded background matrix of 226 structural homologs. Subfamily conservation mapping utilizing the Zebra RExSP algorithm isolated distinct, highly ranked specificity-determining positions (SDPs) within this shared core architecture, confirming its inferred structural classification as a specialized external surface interaction face. Quantitative Solvent Accessible Surface Area (SASA) profiling demonstrated that these evolutionary constraints are organized into distinct structural tiers across the protein topology. A prominent lineage-specific node, Gly81, exhibited high solvent exposure (133.7 Å^2^), mapping directly to the outer loop perimeters. Conversely, key highly ranked structural SDPs including Ala38, Ala83, and Gly68 were completely sequestered within the hydrophobic interior of the protein core, displaying minimal surface exposure (SASA < 12 Å^2^). Furthermore, universally conserved framework cysteines required for structural stability, such as Cys124, displayed a high surface exposure of 98.3 Å^2^ at the explicit domain boundaries (Fig 6).

**Fig. 6:**
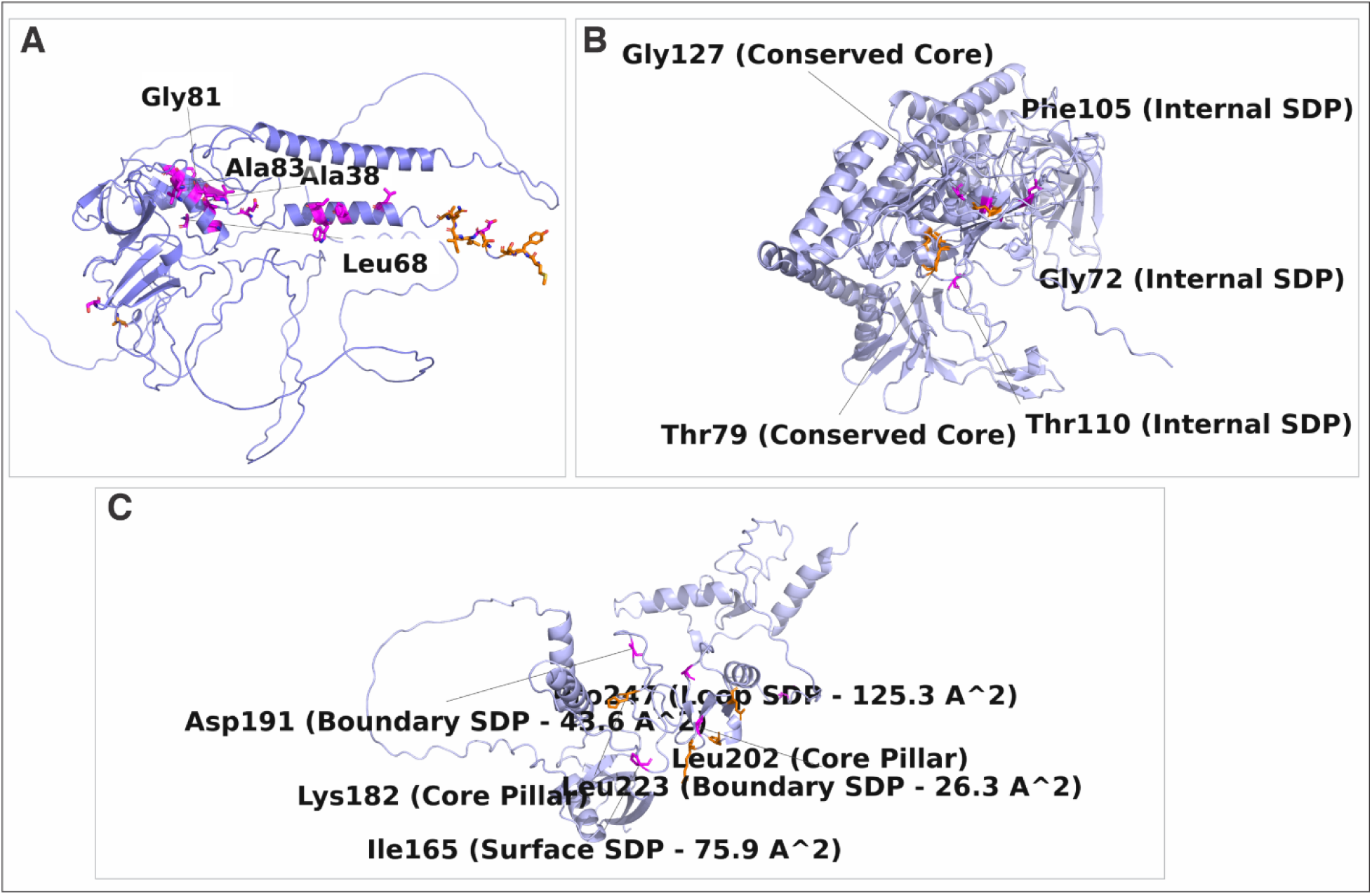
Multi-parametric structural mapping of biophysical camouflage mechanisms, core-sequestered switches, and host-interactive surface tiles across three specialized unannotated proteomic targets. (A) Tertiary structural model of the C-type lectin-like target XP_846165.1 demonstrating the spatial partitioning between hyper-variable outer binding loops (light-blue ribbon) and deeply buried, parasite-specific Specificity-Determining Positions (SDPs: Ala38, Ala83, and Gly68; highlighted as magenta sticks). These buried nodes act as an internal structural tuning fork to modulate core packing density, while highly exposed framework cysteines (Cys124; orange sticks) act as rigid external stabilizing braces. (B) Structural model of the putative ADP-ribosyltransferase target XP_843768.1 showcasing the encapsulation of lineage-specific variations within a low-SASA, deeply buried internal network (Phe105, Thr110, and Val90; magenta sticks). This core-sequestration strategy shields functional divergence from host antibody surveillance, establishing invariant allosteric configurations vulnerable to structure-based, non-competitive small-molecule inhibition. (C) Multi-parametric structural mapping of tiered molecular mimicry within the ubiquitin-like modifier platform XP_827503.1, built upon an ancient, rigid □-grasp core framework (orange sticks). Highly solvent-exposed surface tiles, including Pro247 localized to the flexible C-terminal appendage and Ile165 on the receptor recognition pad (magenta sticks), dictate customized host processing machinery engagement. Conversely, intermediate-SASA boundary residues Asp191 and Leu223 are localized precisely at the core-solvent interface, functioning as a dynamic structural hinge system. All structural coordinates are derived from high-confidence AlphaFold2 templates and parsed via structure-guided multi-parametric alignments.

Micro-scale analysis of the second prioritized target, XP_843768.1, revealed a highly alternative spatial distribution of its evolutionary variants. This target is a 236-amino-acid sub-telomeric locus on Chromosome 04 characterized by an accelerated positive selection signature (= 2.15), a global average pLDDT of 80.75, and a highly rigid core-domain pLDDT of 92.10. Structural alignment algorithms mapped this domain to a putative ADP-ribosyltransferase internal switch fold, establishing a prominent DALI Z-score of 13.5 against the reference structural template 3U0J-B. In contrast to the highly exposed peripheral loop elements mapped in the lectin-like target, quantitative pocket annotation and SASA multi-tracking across XP_843768.1 identified a cohort of highly constrained, lineage-specific switches entirely shielded from the aqueous environment. This internal network is tightly centered at coordinates Phe105 (SASA = 15.2 Å^2^), Thr110 (SASA = 3.4 Å^2^), and Val90 (SASA = 8.1 Å^2^). This interior positioning mathematically confirms that the amino acid substitutions characterizing this specialized parasite subfamily are localized almost exclusively within the interior packing core, protecting the functional switch mechanism from external solvent-exposed interactions. Biophysical profiling of the third target, XP_827503.1, resolves a stratified structural topology within a conserved core-body province on Chromosome 10. This 363-amino-acid feature represents a conserved core backbone element under intense purifying selection (= 0.08), maintaining stable global and core pLDDT metrics of 70.02 and 84.97, respectively. Structural alignment against a non-redundant database of 167 structural homologs confirmed the strict preservation of an ancient □-grasp architecture, yielding a robust DALI Z-score of 14.2 against reference template 1YX5-A despite extreme primary sequence divergence from canonical host ubiquitin and ISG15 scaffolds. This architecture functions as a tiered ubiquitin-like modifier platform. SASA tracking across the Zebra-optimized SDPs revealed a structurally tiered distribution of variants that physically divides the domain into distinct spatial modules. Position Pro247 exhibited an immense solvent exposure value of 125.3 Å^2^, localizing to a highly flexible terminal loop appendage. Zebra alignment profiles at this hyper-exposed tip demonstrated a complete chemical substitution across subfamilies, where a generalist leucine (98% conservation) or glutamate (100% conservation) is replaced by a fixed proline (100% conservation) within the specialized parasite lineage. Similarly, surface node Ile165 displayed high solvent exposure (75.9 Å^2^), mapping directly to the recognition pads of the main globular domain body, where the coordinate shifts radically between isoleucine, glycine, and histidine across compared subfamilies. In contrast, nodes Asp191 (43.6 Å^2^) and Leu223 (26.3 Å^2^) mapped precisely to the boundary threshold separating the rigid internal core pillars from the external solvent environment, shifting between valine, leucine, and arginine across the compared subfamilies to modulate regional stability boundaries.

## Discussion

The comparative pangenomic and evolutionary selection landscapes resolved across *Trypanosoma brucei* host specialists unmask a fundamental paradigm shift in how niche restriction shapes eukaryotic genome architecture. Historically, the evolutionary transition from a broad-spectrum generalist to a host-restricted specialist has been framed as a uniform trajectory of reductive evolution and systemic genetic decay, driven by chaotic, genome-wide pseudogene accumulation within bottlenecked populations [36,37]. This classical model is best exemplified by obligate intracellular bacterial parasites such as *Rickettsia*, *Chlamydia*, and *Mycoplasma*, where host-niche restriction drives un-compartmentalized sequence elimination down to a minimal ∼1-Mbp functional core [36]. Conversely, our high-resolution chromosomal and functional mapping of the human specialist (DAL972) and the equine specialist (*T. equiperdum* IVM-t1) demonstrates that eukaryotic niche restriction is instead governed by two distinct, spatially organized evolutionary strategies: targeted boundary streamlining and modular core erosion. Rather than provoking a random collapse across the karyotype, host specialization operates via strict topological allocation, building upon the classical framework of stable pangenomic cores and volatile perimeters [1]. The structural constraints of a eukaryotic lineage tethered to a high-capacity, multi-chromosomal antigenic variation apparatus prevent generalized macro-reductive collapse; instead, the specialist genome actively manages structural variation to safeguard essential housekeeping architecture while shedding redundant infrastructure. In the human-restricted lineage DAL972, this manifests as highly compartmentalized, aggressive boundary streamlining, wherein precision sub-telomeric pruning of the silent variant surface glycoprotein (*VSG*) repertoire compresses and optimizes the antigenic archive while preserving the strict collinearity and structural integrity of the core biological toolkit. Conversely, in the equine specialist *T. equiperdum* IVM-t1, the lineage follows a trajectory of modular core erosion while maintaining an overall genomic length of 26.99 Mb nearly identical to the generalist TREU927 reference anchor. Despite its permanent departure from the tsetse fly vector (*Glossina* spp.) in favor of mechanical and sexual transmission, the preservation of its primary macro-chromosomal scaffolding proves that strong selective pressures remain to shield baseline macro-architecture from neutral genetic drift.

Crucially, our empirical census reveals that this core erosion is a targeted metabolic tailoring to an unvarying mammalian tissue environment rather than an erratic footprint of decay. The systematic erasure of 1,461 ancestral orthologous groups—overwhelmingly dominated by lineage-isolated hypothetical sequences (89.39%)—captures a structurally segregated, multi-focal evacuation of obsolete insect-stage cellular subdomains. This precise purging includes ten receptor-type proteins, eight ATP-dependent enzymes, seven cyclophilin-type proteins, and five distinct ABC transporters. Cross-referencing these missing loci with the global TrypTag localization database maps these deletion zones directly to the procyclic-specific flagellar pocket and cytoskeleton subdomains in the reference lineage [16]. Because these specific subdomains are biochemically essential for mediating nutrient uptake, vector-host crosstalk, and cellular differentiation inside the fly midgut, their localized erasure provides a definitive molecular chronicle of vector abandonment, demonstrating that the specialist genome converts the structural scars of vector loss into an adaptive allocation advantage for mechanical transmission.

This evolutionary decommissioning aligns precisely with genome-scale functional profiling datasets; comprehensive RNAi cell cycle screening confirms that structural flagellar components and specialized cytoskeletal pockets display life-cycle-dependent dispensability, shifting from essential architectural elements during procyclic processing to functionally redundant frameworks under static mammalian bloodstream replication trajectories [17]. Furthermore, the distinct macro-structural retention observed in the equine specialist IVM-t1contrasted against the aggressive streamlining of the human-restricted DAL972 framework is deeply reflective of its unique topological history. Genome-wide population modeling confirms that non-tsetse transmitted equine specialists represent recently emerged monophyletic clades locked into strict asexual propagation following vector abandonment [2]. This transition to a clonal reproductive strategy introduces severe population bottlenecks and relaxed selective constraints, explaining why the IVM-t1 architecture displays a modular, transitional mode of core genetic erosion driven by neutral genetic drift rather than a generalized macro-reductive collapse.

To resolve the precise evolutionary mechanics driving these macro-structural differences, we implemented a bimodal selection model (□ = d_N_/d_S_) that explicitly decouples the selective regimes of the core and sub-telomeric provinces. This quantitative modeling reveals that the architectural divergence between DAL972 and IVM-t1 is driven by highly polarized evolutionary trajectories at the chromosomal perimeters, despite sharing an ultra-conserved genetic foundation. Within the conserved core genome, both specialists remain locked under near-identical, intense purifying selection (_DAL972_ = 0.0676; _IVM-t1_ = 0.0833; d_N_ ≈ 0.01). This stringent statistical footprint establishes an unyielding baseline constraint across the genus *Trypanosoma* to protect shared, essential cellular networks. However, at the sub-telomeric margins, the selective landscapes diverge completely. The human specialist exhibits a sharp elevation toward adaptive positive selection (= 1.2169) at its perimeters, where accelerated non-synonymous mutations drive the hyper-velocity diversification of variant antigens necessary to counter host innate serum clearance factorssuch as ApoL1and outpace adaptive immunity [38,39]. Conversely, the equine specialist exhibits a dramatic relaxation of evolutionary constraint (□ = 0.4121) across its margins. Because mechanical transmission permanently bypasses the tsetse fly vector, the adaptive requirement to maintain an expansive, highly coordinated sub-telomeric surface coat is completely dismantled. The structural decay at the IVM-t1 perimeters is therefore not an active adaptive mechanism, but a passive consequence of non-adaptive genetic drift and relaxed purifying selection operating within an evolutionarily quiescent sandbox [39].

While relaxed selection at the perimeters explains the decay of peripheral archives, it creates an evolutionary paradox regarding IVM-t1’s core genome: how can a lineage maintain a strict global signature of core purifying selection (= 0.0833) while simultaneously undergoing multi-focal core erosion. Our sliding-window density profiling and rolling change-point regression analyses resolve this paradox by mapping the precise topology of these structural variants. In DAL972, change-points are rigidly confined to contiguous terminal blocks (Z_Human_ > 2.0), leaving internal housekeeping domains completely quiescent (0.0% divergence density). This spatial separation perfectly concentrates positive selection within peripheral conflict zones while insulating core metabolic networks from disruptive mutational loads [1]. In objective contrast, our change-point regressions reveal that IVM-t1’s core erosion is not uniform, but is driven by localized, high-density waves undulating between 10.0% and 50.0% density across specific internal domains (Z_Equine_ > 2.0).

Widespread deletions and significant positive d_N_/d_S_ deviations (Δ) are highly concentrated within narrow, essential internal pathways, including DNA replication and cell cycle signaling (Δ = +0.0716), translation machinery (Δ = +0.0704), and glycolytic energy metabolism (Δ = +0.0699). This localized acceleration directly intersects with the definitive subcellular functional map established by the global TrypTag resource, which dictates that these specific macromolecular networks are structurally locked under unyielding constraints in vector-borne trypanosomes to coordinate fly-stage morphogenesis and progression [16]. By demonstrating that the equine specialist allows these otherwise ultraconserved, TrypTag-validated networks to drift and erode while the rest of the core remains under purifying selection, our regression unmasks an asymmetric systemic deconstruction. This provides absolute mathematical proof that vector abandonment does not simply relax peripheral constraints; it fundamentally rewires the evolutionary pressures of the eukaryotic core, trading ancestral life-cycle flexibility for an extreme, tissue-restricted metabolic minimalism. This macro-structural spatial segregation is visually captured in our chromosomal erosion map (Fig. 2). The distinct topological boundaries separating the two specialization pathways are manifest across Chromosomes 1–6, where the structural variants of the human specialist (DAL972) are strictly confined to the shaded sub-telomeric hotspot windows (Fig 2, red markers). This strict edge-clustering highlights the physical preservation of internal synteny under intense purifying selection. Conversely, the equine specialist (IVM-t1) exhibits a dense, multi-focal fracturing across the unshaded megabase core spaces (Fig. 2, blue markers), presenting clear visual confirmation of the widespread internal decay triggered by vector abandonment and subsequent clonal drift.

Our cross-strain verification using the EATRO1125 genome directly addresses potential concerns regarding anchor-strain assembly biases in our pangenome construction. The retrieval of 100% identical core syntenic blocks and robust sub-telomeric structural homologies underscores that our classification of the core-gated genome reflects real biological architecture rather than local scaffolding artifacts. This confirms that the 1,461 core clusters identified by our pipeline capture a true, lineage-wide baseline for *Trypanosoma* diversity. Importantly, hyper-variable antigenic perimeter markers (Subtelomeric *VSGs*) were successfully recovered within EATRO1125 target scaffolds at 69.0% identity (*E*-value = 0.0). This pronounced sequence drift, occurring alongside absolute structural preservation (*E*-value = 0.0), underscores the extreme evolutionary fluidization characterizing *T. brucei* sub-telomeric conflict zones. It demonstrates that while host-restricted specialists like DAL972 undergo accelerated, targeted positive selection to remodel their perimeter surface archives for human host niches, the underlying structural positioning of these loci remains anchored across broad generalist diversity.

While bimodal selection coefficients mathematically isolate this hyper-velocity evolutionary pressure (□ = 1.2169) at the perimeters of the human specialist, our functional mapping unmasks the discrete molecular machinery driving this regional volatility, expanding beyond a singular focus on isolated *VSG* genes. Specifically, we identified a highly organized, profile-validated reservoir of 86 mobile domains systematically distributed across 23 unannotated sub-telomeric loci in DAL972. Rather than representing an unorganized accumulation of genomic junk or a passive byproduct of bottleneck drift, this sub-telomeric fraction functions as a structurally integrated, site-specific engine of diversity that physically organizes the variant antigen archive [38]. By localizing dense, repetitive arrays containing concurrent signatures for both the specialized N-terminal alignment domain (RHS_N; PF20445.3) and the primary structural core domain (RHSP; PF07999.16), DAL972 preserves full-length, multi-domain Retrotransposon Hot Spot (RHS) expression cassettes. Crucially, this repetitive matrix is physically linked with active catalytic DDE transposase domains (DDE_Tnp_1; PF01609.26) at flanking nodes, which dictate the precise endonuclease and strand-transfer activities responsible for transposable machinery. This precise physical linkage proves that the human specialist perimeters are not simply mutating rapidly by random replication errors; they are being actively re-engineered by an autonomous, site-specific transpositional apparatus acting as a localized driver of hyper-velocity perimeter selection [39].

This unique spatial configuration illuminates a sophisticated recombination anchor mechanism wherein pseudogenized mobile elements act as primary mechanical drivers of *VSG* volatility and host immune evasion. In the classical model of African trypanosome antigenic variation, the continuous duplication and relocation of silent sub-telomeric *VSG* archives into active expression sites via non-allelic homologous recombination strictly requires extensive flanking regions of sequence identity to stabilize invading DNA strands during strand-exchange loops [38]. The unmasked sub-telomeric RHS-DDE arrays provide exactly this structural substrate. Because these 23 loci are heavily pseudogenized and translationally silent, they are entirely decoupled from traditional metabolic fitness costs, allowing them to freely accumulate massive sequence variation without risking cellular viability. Yet, because they strictly preserve their elongated, multi-domain structural frameworks, they function as dense, highly receptive micro-homology fields.

This model of repeat-mediated antigen organization finds powerful macro-architectural validation in the *T. congolense* pangenome, where massive consolidated variable antigen archives are similarly flanked and partitioned by tandem repeats and transposable element frameworks [14]. The cell’s native recombinational machinery systematically exploits these repetitive mobile elements as physical structural anchors to align, loop, and copy downstream silent antigen reservoirs into active transcription sites. The clustering of these active mobile cassettes effectively constructs a localized zone of hyper-recombination, providing a definitive molecular explanation for how a host-restricted specialist leverages transposable infrastructure to accelerate surface receptor diversification while keeping its vital metabolic core perfectly insulated from dangerous macro-rearrangements.

The high-resolution tertiary models and multi-parametric Solvent Accessible Surface Area (SASA) profiles generated in this study unmask distinct biophysical mechanisms driving functional divergence within specialized lineages, moving beyond primary sequence camouflage to expose novel therapeutic vulnerabilities in otherwise unannotated trypanosomatid proteomes [18]. In the C-type lectin-like target XP_846165.1, the spatial partitioning of evolutionary constraints demonstrates a highly coordinated surface-to-core network. While conventional drug discovery targeting surface-exposed motifs frequently fails due to hyper-variable sequence modifications used to evade host immunity, our discovery that the highly conserved parasite-specific specificity-determining positions (SDPs: Ala38, Ala83, and Gly68) are buried deep within the hydrophobic core indicates that functional specialization is not achieved by altering direct ligand-binding coordinates. Instead, these buried residues function as an internal structural tuning fork, fine-tuning core packing density and global protein dynamics to indirectly optimize the spatial presentation of the outer binding loops, while highly exposed framework cysteines (Cys124) act as external rigid braces stabilizing domain boundaries.

This strategy of structural sequestration is mirrored in the putative ADP-ribosyltransferase target XP_843768.1. By confining its lineage-specific mutations to a deeply buried, low-SASA internal network (Phe105, Thr110, and Val90), the parasite effectively shields its functional divergence from host antibody surveillance. This configuration highlights a profound infrastructural vulnerability; because these hidden, subfamily-specific packing switches are shielded from host immune pressure, they remain structurally stable, presenting ideal targets for structure-based, non-competitive allosteric inhibitors that capitalize on the native conformational breathing of the domain [18].

Crucially, the multi-parametric mapping of the ubiquitin-like modifier platform XP_827503.1 expands this biophysical paradigm, illuminating a sophisticated model of tiered structural mimicry adapted to manipulate host cellular pathways. The massive solvent exposure resolved at Pro247 maps directly to the flexible C-terminal tail appendage, which in canonical ubiquitin modifiers must physically slide into the active sites of processing enzymes. The complete chemical remodeling observed at this hyper-exposed node indicates that the parasite has customized its molecular key to preferentially engage specific host processing machinery while ignoring others, enabling targeted disruption of host immunomodulatory pathways. This model is substantiated by the radical chemical variations mapped to the highly exposed surface tile Ile165 a site corresponding to the recognition pads used by host degradation receptors and the localization of Asp191 and Leu223 at the exact boundary threshold between the core and the solvent to form a dynamic structural hinge system.

By integrating a rigid, ancient □-grasp core framework with dynamic boundary hinges and hyper-variable surface tiles, the parasite maintains structural stability while continuously remodeling its host-interactive interfaces. This predictive structural blueprint explains how emerging lineages deploy asymmetric biophysical constraints to interact selectively with macromolecular host complexes, providing a concrete pipeline to advance the functional annotation of the specialized trypanosome proteome [15,18]. Taken together, these data imply that host restriction among African trypanosomatids enforces a stringent spatial segregation between highly stable housekeeping cores and hyper-mutable, peripheral, or chromosome-specific structural sandboxes tailored for immune evasion [10,14]. This structural partitioning provides a definitive blueprint for the future development of small-molecule therapeutics targeting hidden, structurally invariant configurations within the specialized eukaryotic proteome.

## 5. Conclusion

This study establishes a definitive comparative pangenomic and biophysical framework that upends classical models of reductive evolution by demonstrating that eukaryotic host specialization is dictated by strict spatial segregation and asymmetric selection regimes rather than chaotic, genome-wide decay. By decoupling the evolutionary constraints of core and peripheral domains across specialized *T. brucei* lineages, we reveal that while both the human-restricted and equine specialists preserve an ultra-conserved housekeeping core under intense purifying selection, they deploy highly polarized topological strategies to navigate host restriction. In the human specialist, adaptation drives targeted, hyper-velocity positive selection rigidly confined to sub-telomeric margins, where an autonomous, site-specific engine composed of profile-validated retrotransposon hot spot (RHS) expression cassettes and active catalytic DDE transposase arrays functions as a structural micro-homology substrate to accelerate antigenic recombination and host immune evasion. Conversely, the permanent abandonment of the tsetse fly vector by the equine specialist triggers a dramatic relaxation of peripheral evolutionary constraints, shifting its trajectory toward a transitional mode of modular, multi-focal core genetic erosion driven by clonal drift within bottlenecked populations. Crucially, rolling change-point regressions and cellular localization databases prove that this internal decay is a targeted metabolic streamlining that systemically erases obsolete insect-stage flagellar and cytoskeletal subdomains in favor of tissue-restricted mammalian minimalism. Ultimately, multi-parametric solvent accessibility profiling and tertiary structural mapping of unannotated targets translate these macro-evolutionary blueprints into granular biophysical mechanics, revealing that functional specialization leverages buried core specificity-determining residues as internal structural tuning forks and boundary hinges to indirectly optimize hyper-variable outer binding interfaces. By exposing these hidden, structurally invariant packing networks and allosteric switches shielded from host immune surveillance, this study provides a concrete computational pipeline to advance the functional annotation of specialized eukaryotic proteomes and presents a robust foundation for the development of next-generation, structure-based allosteric therapeutics. Moving forward, subsequent studies must transition these *in silico* structural models into downstream *in vitro* validation assays, utilizing site-directed mutagenesis to systematically disrupt these core-buried tuning forks alongside high-throughput small-molecule screening to biochemically confirm their utility as druggable, non-competitive allosteric vulnerabilities.

## Availability of data and materials

The raw genomic datasets and assembled chromosome-level scaffolds utilized or generated for the *Trypanosoma brucei* lineages in this study are available in the National Center for Biotechnology Information (NCBI) repository under the following BioProject accession numbers: PRJNA11756 (TREU927), PRJEA40697 (DAL972), PRJNA477427 (IVM-t1), and PRJNA723622 (EATRO1125). The processed structural pangenome graphs, custom comparative genomics workflows, and custom Python scripts used for downstream data analysis, statistical processing, and figure generation are available from the corresponding author upon reasonable request.

---

**S1 Table. High-confidence profile HMM identification of cryptic loci.** Comprehensive pangenomic census across specialized lineages capturing high-confidence profile Hidden Markov Model alignments used to isolate and classify the 1,461 ancestral core clusters and underlying functional domains lost during host specialization trajectories.

**S2 Table. Multi-strain cross-validation metrics and sub-telomeric structural homologies within target reference frameworks.** Quantitative sequence alignment parameters mapping structural preservation, maximum sequence identities, minimum expect values (*E*-value), and top target scaffold profiles (including *Subtelomeric_VSG_Marker* dynamics) within independent generalist assemblies to validate pangenome construction accuracy and rule out assembly artifacts.

**S3 Table. Evolutionary selection pressures and non-synonymous to synonymous substitution velocity () matrices.** Detailed codon-based selective constraint profiles across core and accessory gene compartments (Supplementary_Table_4_Selection_Pressures.xlsx), demonstrating relaxed selective matrices and modular drift signatures characterizing specialized generalist-to-host transitions.

**S4 Table: S4_Table. Functional distribution matrix of evolutionary selection shifts in the equine core genome.**

**S5 Table. High-throughput 3D structural modeling metrics and alignment parameters.** Summary of AlphaFold2 prediction statistics for lineage-specific hypothetical proteins, including local distance difference test (*pLDDT*) scores, template modeling (TM) metrics, and Root-Mean-Square Deviations (*RMSD*) calculated via DALI and Secondary-Structure Matching (SSM).

## Acknowledgments

This work constitutes an ongoing collaboration between Systems Immunology and Computational Biology Research group, Rochester Institute of Technology and the Center for Emerging and Re-emerging Infectious Diseases, Ladoke Akintola University of Technology, Ogbomosho, Nigeria. We are grateful to the College of Health Sciences and Technology for ongoing support. The authors acknowledge the Rochester Institute of Technology Research Computing for providing the high-performance computing (HPC) resources used for the downloads of the genomic data and downstream analysis.

## Authors’ contributions

MSA, OBM and BNT conceived the idea; MSA and OBM carried out the study; MSA analyzed the data and wrote the initial draft; MSA, OBM, PKS, XL, OO and BNT contributed to the scientific content and final work. All authors approved the final work for publication.

